# Human sensory Long-Term Potentiation (LTP) predicts visual memory performance and is modulated by the brain-derived neurotrophic factor (*BDNF*) Val^66^Met polymorphism

**DOI:** 10.1101/284315

**Authors:** M.J. Spriggs, C.S. Thompson, D Moreau, N.A. McNair, C.C. Wu, Y.N. Lamb, N.S. McKay, R.O.C. King, U. Antia, A.N. Shelling, J.P. Hamm, T.J. Teyler, B.R. Russell, K.W. Waldie, I.J. Kirk

**Author notes:** M.J.S and C.S.T contributed equally to this work. Corresponding author: Meg J. Spriggs, School of Psychology, University of Auckland. Private Bag 92019, Auckland 1142, New Zealand.

## Abstract

**Background:** Long-Term Potentiation (LTP) is recognised as a core neuronal process underlying long-term memory. However, a direct relationship between LTP and human memory performance is yet to be demonstrated. The first aim of the current study was thus to assess the relationship between LTP and human long-term memory performance. With this also comes an opportunity to explore factors thought to mediate the relationship between LTP and long-term memory, and to gain additional insight into variations in memory function and memory decline. The second aim of the current study was to explore the relationship between LTP and memory in groups differing with respect to *BDNF* Val^66^Met; a single nucleotide polymorphism implicated in memory function.

**Methods:** 28 participants (15 female) were split into three genotype groups (Val/Val, Val/Met, Met/Met) and were presented with both an EEG paradigm for inducing LTP-*like* enhancements of the visually-evoked response, and a test of visual memory.

**Results:** The magnitude of LTP 40 minutes after induction was predictive of long-term memory performance. Additionally, the *BDNF* Met allele was associated with both reduced LTP and reduced memory performance.

**Conclusions:** The current study not only presents the first evidence for a relationship between sensory LTP and human memory performance, but also demonstrates how targeting this relationship can provide insight into factors implicated in variation in human memory performance. It is anticipated that this will be of utility to future clinical studies of disrupted memory function.

## Introduction

First demonstrated *in vivo* in 1973 ^1^, Long-Term Potentiation (LTP) has since been widely recognized as the principal model for the neuronal basis of long-term memory. LTP is an enduring facilitation of synaptic transmission between neurons that follows repeated co-activation of the neurons in a network ^2–4^. The cellular and molecular mechanisms of LTP have been studied extensively *in vivo* and *in vitro* in laboratory animals, which typically involves the application of direct neuronal electrical stimulation and results in an enhancement of the response in a neighbouring cell ^1, 4–7^. Such studies have demonstrated that, in its most widespread form, LTP is dependent on the influx of Ca^2+^ through N-methyl-D-aspartate (NMDA) receptors, leading to long-term alterations in cell structure and function, and an increase in synaptic efficacy. However, a direct relationship between LTP and memory performance has been notoriously difficult to demonstrate. *In vivo* rodent studies have typically focused on spatial memory as a measure of memory function ^8^. While this is an accessible measure of memory, the disparate results from such studies indicate that it may not be the most appropriate index of LTP. As such, a definitive demonstration of the relationship between LTP and memory performance remains elusive.

Using similar induction protocols, the properties of human LTP have been shown to be consistent with those seen in animals^9, 10^. However, due to the invasive nature of these procedures, such studies have been limited to excised human tissue. The sensory LTP paradigm presents one of the first opportunities for the non-invasive *in vivo* study of an LTP-*like* shift in Event Related Potentials (ERPs) in humans. First presented by Teyler et al., ^11^, the sensory LTP paradigm typically involves presenting participants with high-frequency (∼9Hz) visual stimulation, which leads to an enhancement of the visually-evoked potential (VEP) to subsequent low-frequency (∼1Hz) presentations of the same stimulus. This enhancement has been shown to conform to many of the Hebbian characteristics of LTP ^12–14^, and is generated by a bottom-up modulation of connection strength between occipital and temporal regions ^15^. As such, this experience-dependent enhancement of the VEP is understood to represent the induction of an LTP*-like* form of neuroplasticity ^16, 17^.

As the sensory LTP paradigm can be used non-invasively with humans, it provides a novel avenue for assessing the pivotal relationship between LTP and human long-term memory performance. The primary aim of the current study was thus to assess this relationship using the visual LTP paradigm and two subtests of the Wechsler Memory Scale-III ^18^ that are widely used in clinical assessments of delayed visual recognition memory^19–22^. It was hypothesized that individuals with greater LTP magnitude would also demonstrate greater memory performance.

With the ability to study LTP and memory performance comes an unique opportunity to also study the role that LTP alterations play in variations in human memory performance. Previous studies using the human sensory LTP paradigm have demonstrated modulated LTP in healthy populations differing in physical fitness ^23^, age ^24, 25^, and genetics ^15^, as well as in clinical conditions such as depression ^26^ and schizophrenia ^27^. However, how these differences in LTP magnitude impact on memory performance is yet to be assessed.

One factor implicated in healthy variations in human memory performance is the gene that controls the secretion of brain-derived neurotrophic factor (BDNF). BDNF regulates neuronal proliferation and differentiation in the developing brain, and is an important molecular mediator of synaptic plasticity in the mature brain ^28–30^. In humans, approximately 25-50% of the population ^31^ carry a single nucleotide polymorphism (SNP) on the *BDNF* gene, which substitutes valine to methionine at codon 66 (Val^66^Met; SNP rs6265). The Met allele of the polymorphism has been associated with reduced declarative memory performance ^32^. However, it is unclear whether these genotype differences are due to the regulatory role of BDNF in brain development ^33^ or due to the modulation of synaptic plasticity ^15, 34, 35^. The secondary aim of this study was thus to explore the effect of the polymorphism on both visual LTP and visual long-term memory. We hypothesized a consistent genotype difference across the two measures (LTP and memory), which would support the notion that the polymorphism mediates long-term memory performance through an effect on LTP.

## Methods and Materials

### Subjects

Fifteen females (all right-handed) and thirteen males (3 left-handed) with a mean age of 24.2 years (range 21-35; *SD* = 3.3 years) took part in this experiment. All participants had normal or corrected-to-normal vision. Subjects gave their informed consent to participate in the study and all experimental procedures were approved by the University of Auckland Human Participants Ethics Committee.

### BDNF Genotyping

DNA was extracted from blood samples using the method described in previous literature ^36^, and were analysed by the Auckland Sequenom Facility. Amplification was carried out on the 113 bp polymorphic BDNF fragment using a polymerase chain reaction (PCR), with Taq polymerase and the following primers: BDNF-F 5’-GAG GCT TGC CAT CAT TGG CT-3’ and BDNF-R 5’-CGT GTA CAA GTC TGC GTC CT-3’. PCR conditions were as follows: denaturation at 95°C for 15 min, 30 cycles on a thermocycler (denaturation at 94°C for 30sec, annealing at 60°C for 30sec, and extension at 72°C for 30sec) with a final extension at 72°C. The PCR product (6.5µL) was incubated with PmlI at 37°C overnight and digestion products were analysed using a High-res agarose gel (4%) with a Quick load 100bp ladder (BioLabs) and a GelPilot Loading Dye (QIAGEN). Digestion resulted in a 113 bp fragment for the Met^66^ allele and this was cut into 78 and 35 bp fragments for the Val^66^ allele. Subjects were divided into three groups defined by Val^66^Met genotype (10 Val/Val, 10 Val/Met and 8 Met/Met).

### Memory Performance

Memory performance was assessed using two subtests of the Wechsler Memory Scale-III (WMS-III^18^: the Faces task, and the Family pictures task.

The Faces task involved presenting participants with a set of 24 faces for two seconds each that they were asked to remember. After a 30-minute delay, participants were tasked with identifying the original faces (make an “old/new” decision) from a selection of 48 faces which included the 24 original faces as well as 24 new faces.

For the Family Pictures task, participants were presented with pictures of a family in a variety of scenes for 10 seconds per scene. Again, after a 30 minute delay, participants were asked to recall details about each scene.

For both tasks, performance was scored as percentage correct. An average score for the two tasks was then used as an index of visual-memory performance.

### Apparatus

EEG data were recorded using a 128-channel Ag/AgCl electrode net (Electrode Geodesics Inc., Eugene, OR) at a continuous sampling rate of 1000Hz (0.1-100Hz analogue bandpass filter) with impedance kept below 40kΩ. EEG data were recorded using a common vertex reference (Cz) and later re-referenced to average offline. Stimuli were presented on an SVGA computer monitor (1024×768 pixel resolution; 60Hz refresh rate).

### Stimuli

Stimuli consisted of two circles containing black and white sine gratings of horizontal and vertical orientation with a spatial frequency of one cycle per degree (size: 9.6 x 9.6cm, 272 x 272 pixels; Figure 1a). Stimuli were presented in full contrast against a grey background in the centre of the screen, subtending a diameter of 8° of visual angle. A red fixation dot was present throughout testing. Stimulus presentation was controlled using E-prime v1.1 (Psychology Software Tools).

**Figure 1.**
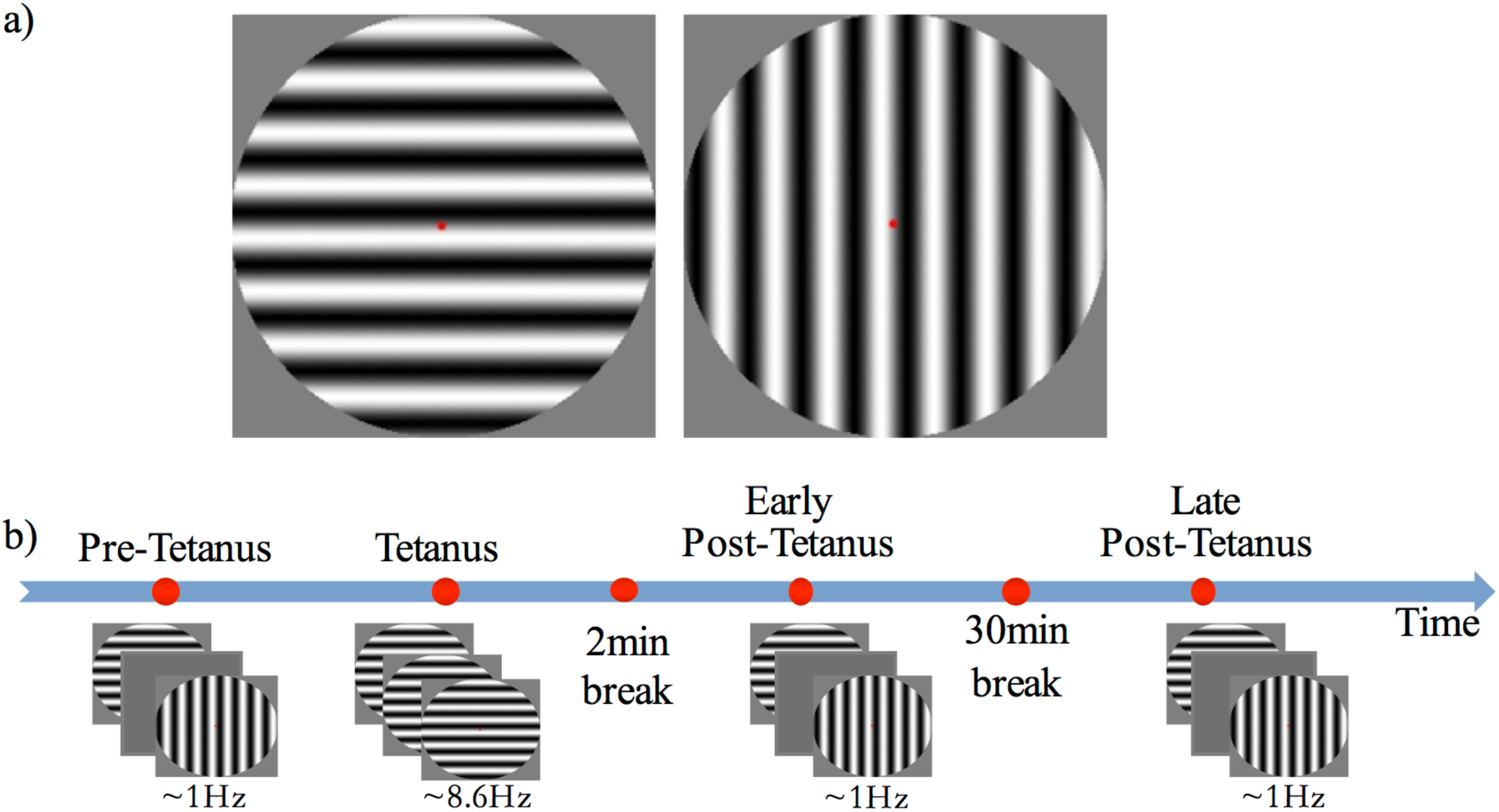
a) The two circular sine gratings of vertical and horizontal orientation used for visual stimuli. b) The experimental procedure consisted of four experimental blocks: three low-frequency blocks (pre-tetanus, early post-tetanus, late post-tetanus), and one high frequency tetanus. Magnitude of LTP was determined by subtracting pre-tetanus amplitude from each of the post-tetanus blocks. Rest periods were included after the tetanus (2 minutes), and between the two post-tetanus blocks (30 minutes).

### LTP procedure

LTP was assessed using our previously established EEG paradigm ^11, 13, 14, 17, 23^ (Figure 1b). Participants were first presented with a ‘pre-tetanus’ baseline block, consisting of 240 presentations of each stimulus at a low temporal frequency of 0.67-1Hz (33ms presentation with a jittered inter-stimulus interval of 1000-1500ms, ∼8 minutes). Each participant was then presented with one of the stimulus orientations (counterbalanced) as an LTP-inducing stimulus, or ‘visual tetanus’. This consisted of 1000 presentations, at a frequency of 8.6Hz (jittered ISI of 67-100ms, ∼2 minutes). This was immediately followed by a two minute rest period to allow retinal afterimages to dissipate. Participants were then presented with two more experimental blocks: an ‘early post-tetanus’ block, and a ‘late post tetanus block’. Both post-tetanus blocks had identical parameters to the pre-tetanus block (240 presentations of each stimulus at a temporal frequency of 0.67-1Hz). The post-tetanus blocks were separated by a 30-minute eyes closed rest period.

### EEG analysis

EEG data were processed using in-house software. The data were segmented into 600ms epochs (−100 – 500ms), band-pass filtered (0.10-30Hz, bidirectional three-pole Butterworth filter ^37^) and baselined to the pre-stimulus period. Epochs containing significant artefacts (e.g., eye-blinks) were removed, and the remaining data were averaged according to block (pre-tetanus, early post-tetanus, and late post-tetanus) and stimulus condition (tetanized, non-tetanized). The magnitude of LTP was defined as the amplitude difference between the pre-tetanus baseline and the two post tetanus blocks independently (referred to as *early LTP* and *late LTP*, respectively) within the N1b time window. In accordance with previous literature^13^, the N1b was defined as the section of the VEP extending from the peak of the N1 to the midpoint between the peak of the N1 and the peak of the P2. The pre-and post-N1b components of the VEP were averaged across a posterior clusters of electrodes that was determined from the topography of the mean visually-evoked potential across all conditions and was centred approximately around P7 and P8 for each participant.

### Statistical analyses

We used Bayesian hypothesis testing for all analyses. Because we understand readers may wish to compare these with frequentist analyses, we provide all the frequentist equivalents in a subsequent section. All analyses were performed in R ^38^, with the following packages: BayesFactor ^39^, dplyr ^40^, ggplot2 ^41^, rjags ^42^.

For all Bayesian analyses, Markov chain Monte Carlo methods were used to generate posterior samples via the Metropolis-Hastings algorithm. All analyses were set at 10,000 iterations, with diagnostic checks for convergence. One chain per analysis was used for all analyses reported in the paper, with a thinning interval of 1 (i.e., no iteration was discarded). All priors used in the reported analyses are default prior scales ^39^.

## Results

### Bayesian analyses

A linear regression analysis with a default prior (r scale of 0.354) showed that *late LTP* was a reliable predictor of *Memory performance* [*P*(M|data) = .77, BF_M_ = 3.27]. The correlation between *LTP* and *Memory performance* was robust (*r* =.44, BF_10_ = 3.32). The association between the two variables, with the breakdown into genotype groups, is shown in Figure 2. In contrast, the data showed no evidence for *early LTP* being a reliable predictor of *Memory performance* [*P*(M|data) = .26, BF_M_ = 0.35], substantiated by a null correlation (*r* =.02, BF_10_ = 0.24).

**Figure 2.**
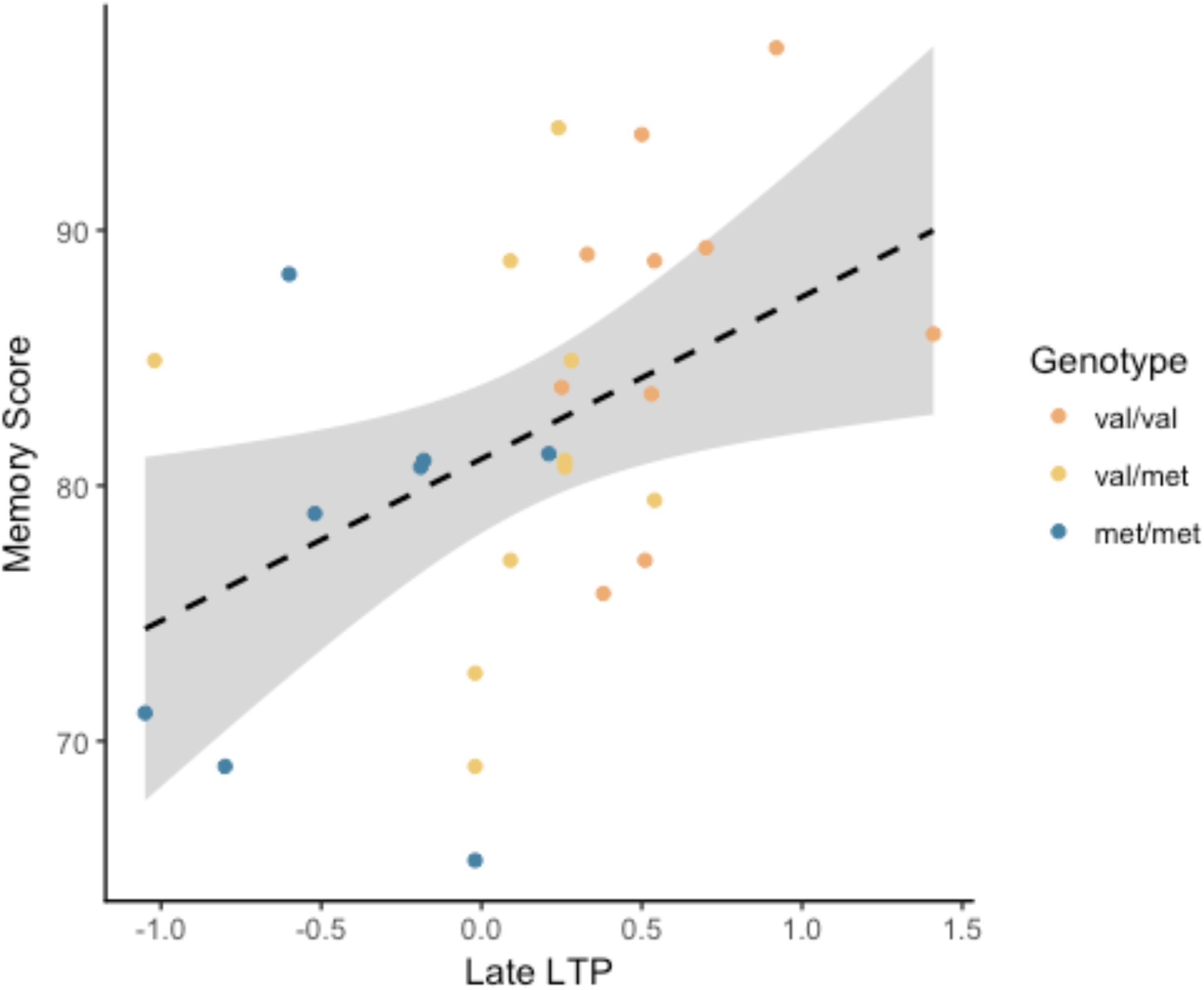
Late LTP amplitude plotted against WMS-III visual memory index (Memory), broken down by genotype. The dashed line is the regression line from the linear model, for all genotype groups combined. The shaded area around the dashed line represents the 95% confidence interval for predictions from the linear model fitted.

An ANOVA with a default prior (r scale of 0.5) on *late LTP* with *BDNF genotype* as a fixed factor, showed overwhelming evidence for the alternative model over the null model [*P*(M|data) > .99, BF_M_ = 338.1, see Fig. 3]. Pairwise comparisons with default prior (r scale of 0.701) showed support, although of different magnitudes, for all possible comparisons. There was overwhelming evidence for the difference between Val/Val and Met/Met (BF_10_ = 407.56), strong evidence for the difference between Val/Met and Val/Val (BF_10_ = 8.06), and very moderate evidence for the difference between Val/Met and Met/Met (BF_10_ = 2.26).

**Figure 3.**
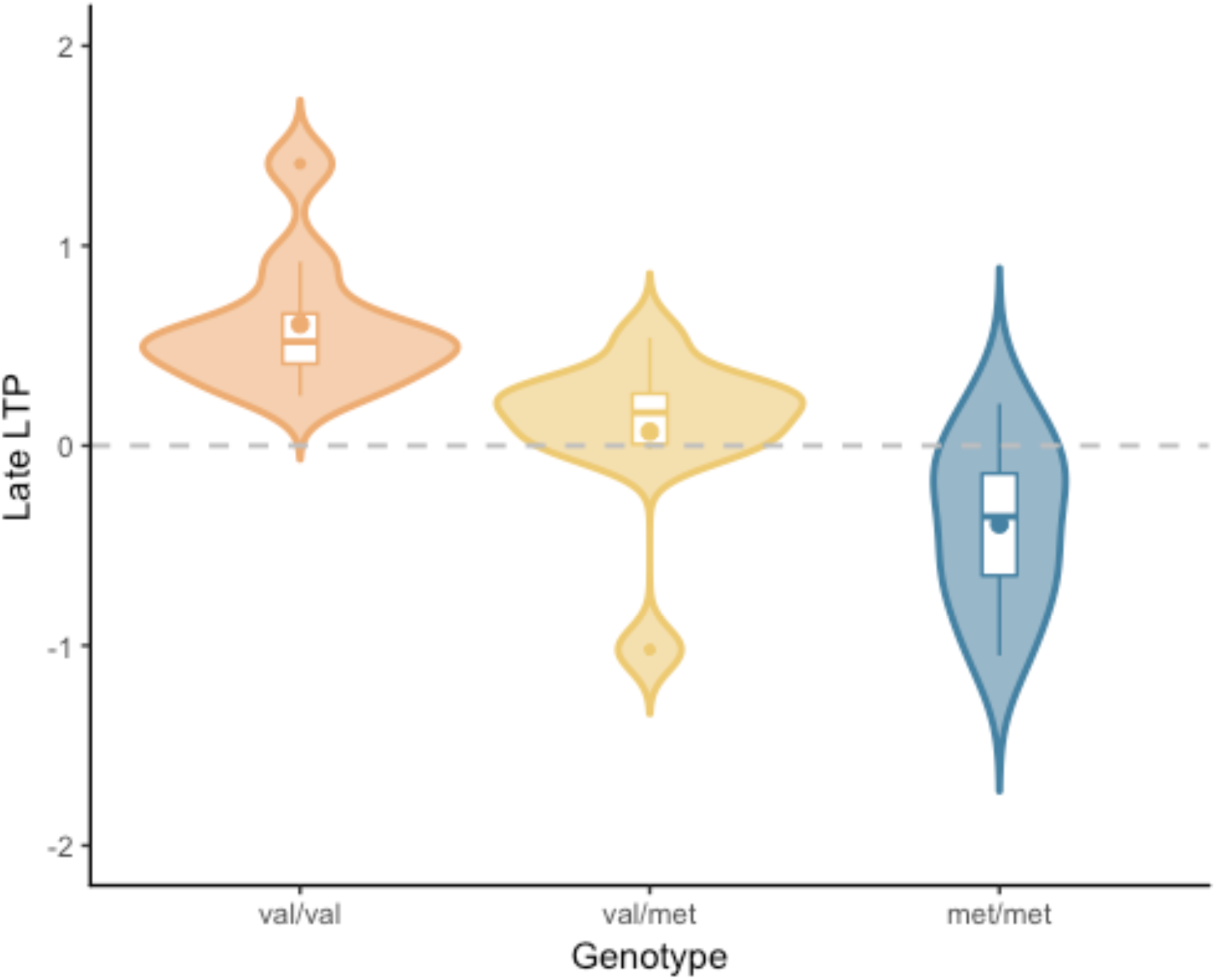
Late LTP amplitude as a function of BDNF genotype. The plot shows the distribution of gain scores (violin) together with the mean (box central dot), median (box central line), first and third quartile (box edges), minimum and maximum (whiskers), and outliers (outside dots).

An ANOVA with a default prior (r scale of 0.5) on *early LTP* with *BDNF genotype* as a fixed factor showed evidence for the alternative model over the null model [*P*(M|data) = .86, BF_M_ = 6.13, see Fig. 4]. Pairwise comparisons with default prior (r scale of 0.701) showed that the effect differed based on the specific comparison. There was moderate evidence for the difference between Val/Val and Met/Met (BF_10_ = 3.18), somewhat stronger evidence for the difference between Val/Met and Val/Val (BF_10_ = 5.46), and no evidence for the difference between Val/Met and Met/Met (BF_10_ = 0.41).

**Figure 4.**
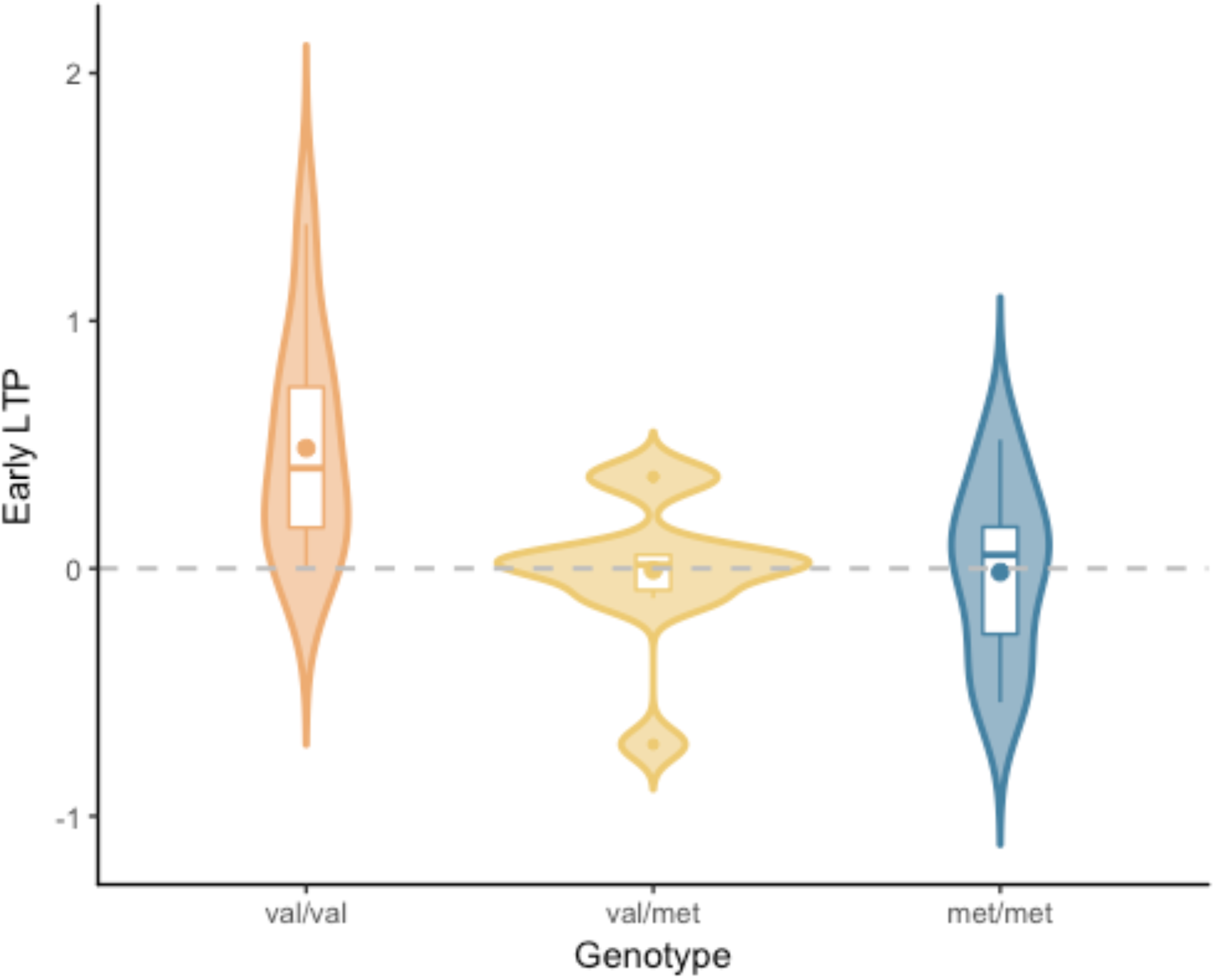
Early LTP amplitude as a function of BDNF genotype. The plot shows the distribution of gain scores (violin) together with the mean (box central dot), median (box central line), first and third quartile (box edges), minimum and maximum (whiskers), and outliers (outside dots).

An ANOVA with a default prior (r scale of 0.5) on *Memory performance* with *BDNF genotype* as a fixed factor, showed moderate evidence for the alternative model over the null model [*P*(M|data) = .69, BF_M_ = 2.19, see Fig. 5]. Pairwise comparisons with default prior (r scale of 0.701) showed support only for the difference between Val/Val and Met/Met (BF_10_ = 4.31). There was no evidence for a difference between Val/Met and Val/Val, or between Val/Met and Met/Met (BF_10_ = 0.99 and BF_10_ = 0.67, respectively).

**Figure 5.**
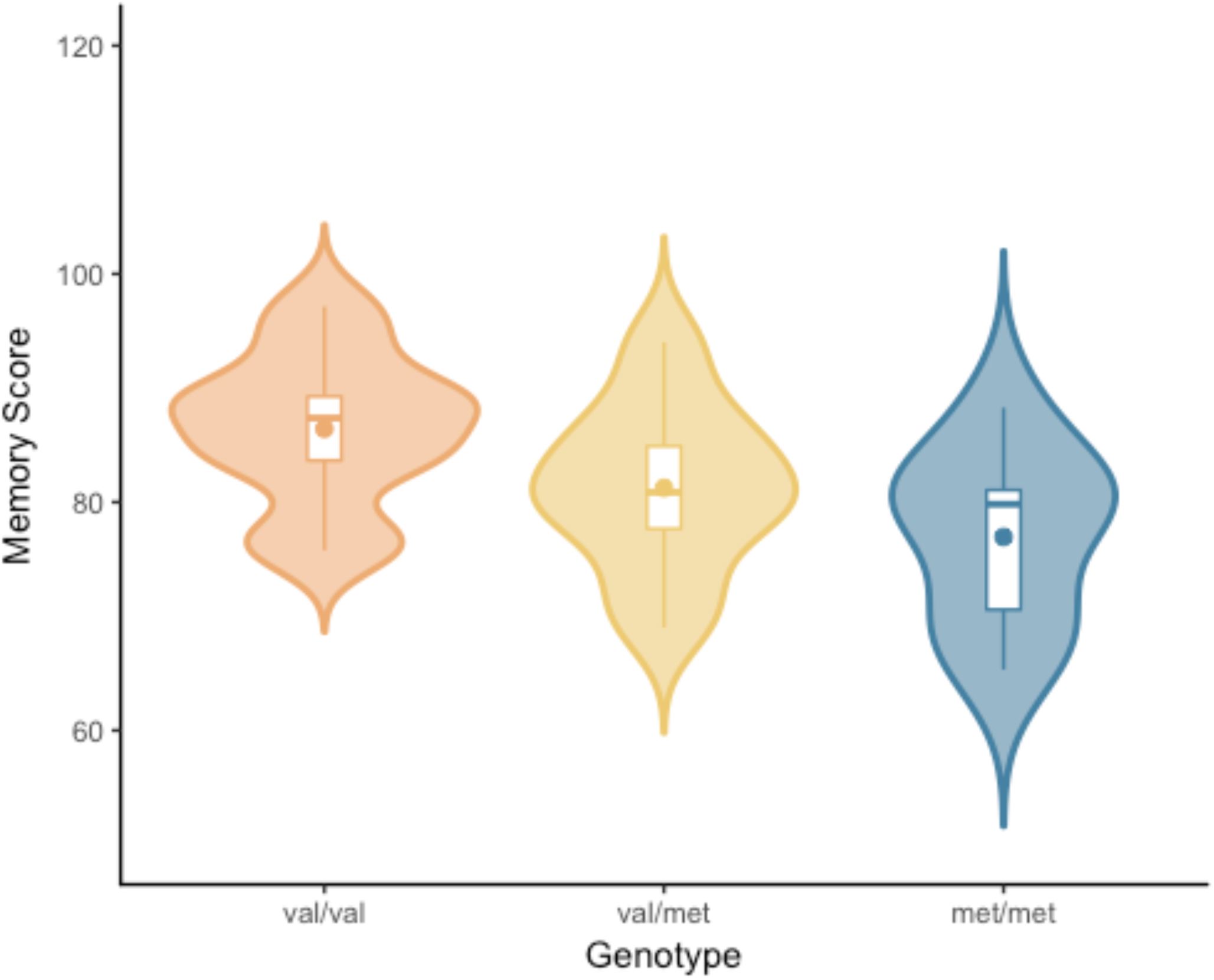
Memory performance as a function of BDNF genotype. The plot shows the distribution of gain scores (violin) together with the mean (box central dot), median (box central line), first and third quartile (box edges), minimum and maximum (whiskers), and outliers (outside dots).

### Frequentist analyses

A linear regression analysis showed that *late LTP* was a reliable predictor of *Memory performance* (*R^2^* = .196, *R^2^_adj_* = .165, *RMSE* = 7.28). The correlation between *LTP* and *Memory performance* was robust [*r* =.44, 95CI = (.08, .70), *p* = .02]. In contrast, *early LTP* was not a significant predictor of *Memory performance* (*R^2^* = .00, *R^2^_adj_* = -.038, *RMSE* = 8.12), and the correlation between the two variables was null (*r* =.02, *p* = .92).

An ANOVA on *LTP* with *BDNF genotype* as a fixed factor showed a significant effect of *BDNF genotype* [*F*(2,25) = 14.61, *p* < .001, see Fig. 3]. Pairwise comparisons showed a significant difference between Val/Val and Met/Met (*p* < .001), between Val/Met and Val/Val (*p* = .006), and between Val/Met and Met/Met (*p* = .03). Note that the latter would not survive correction (FDR or Bonferroni), but all effects are presented uncorrected herein for comparative purpose with the Bayesian analyses.

An ANOVA on *early LTP* with *BDNF genotype* as a fixed factor showed a significant effect of *BDNF genotype* [*F*(2,25) = 5.68, *p* = .009, see Fig. 4]. Pairwise comparisons showed a significant difference between Val/Val and Met/Met (*p* = .02), between Val/Met and Val/Val (*p* = .02), but not between Val/Met and Met/Met (*p* >.999).

An ANOVA on *Memory performance* with *BDNF genotype* as a fixed factor, showed a significant effect of *BDNF genotype* [*F*(2,25) = 3.87, *p* = .03, see Fig. 5]. Pairwise comparisons showed a significant difference between Val/Val and Met/Met (*p* = .02), but not between Val/Met and Val/Val, or between Val/Met and Met/Met (*p* = .12 and *p* = .25, respectively). All effects are presented uncorrected herein for comparative purpose with the Bayesian analyses.

## Discussion

The current study provides the first evidence that the degree of visually-induced LTP is a significant predictor of human visual memory performance. While LTP is at the core of our understanding of long-term memory formation, a direct relationship between the extent of LTP and memory task performance has been notoriously difficult to demonstrate. However, the visual LTP paradigm presents the unique opportunity to study LTP in humans non-invasively, thus allowing for a direct assessment of this paramount relationship.

The current results demonstrate that LTP magnitude in the late post-tetanus block is a reliable predictor of long-term memory performance. This late post-tetanus block is run approximately 40 minutes after the LTP-inducing visual tetanus, and thus indexes enduring changes in neuronal activation. Conversely, there was no relationship between memory performance and early post-tetanus LTP, which indexes immediate post-tetanus modulations in neuronal response. Although these results may appear somewhat contradictory, this indicates that the two post-tetanus blocks index different phases of LTP: LTP induction and LTP maintenance. LTP induction and maintenance are dependent on different cellular processes ^2, 43^, and therefore it is unsurprising that they have distinct relationships to long-term memory performance. Importantly, the current results demonstrate that it is the late post-tetanus block, and thus LTP maintenance, which is key to memory performance, with greater LTP magnitude predicting better memory performance.

The two subtests of the WMS-III assessed here are widely used to evaluate delayed visual recognition memory across diverse clinical populations ^19–22^. Although delayed recognition memory is traditionally thought of as dependent on a network within the medial temporal lobe and frontal cortex^44^, The current results indicate that LTP measured over the visual cortex is predictive of performance on this task. This could therefore be interpreted as representing a general propensity for an individual to exhibit LTP, perhaps across the brain as a whole. However, previous studies of the *BDNF* polymorphism indicate that there is divergence of the effect of the polymorphism between the motor cortex ^45–48^ and auditory cortex ^48^. This suggests that the propensity for plasticity is not homogeneous across the brain, and therefore, sensory LTP is not a global index.

An alternative, and perhaps more likely, explanation for the correlation between visual cortical plasticity and memory performance is the integral involvement of visual system circuitry in networks specifically sub serving visual memory formation. Experience-dependent plasticity within the visual network is understood to be fundamental in the mnemonic processing of visual information ^49–52^, and plastic processes in the visual system influence subsequent processing in hippocampus ^52^. In support of this, Spriggs, Sumner et al., ^15^ recently demonstrated that LTP induction using the visual paradigm modulates connections between the occipital and temporal cortices. It therefore appears that visually-induced LTP may represent an early, yet integral stage in visual memory processing, and that the magnitude of LTP can act as a ‘window’ into the efficacy of the visual memory network.

Consistent with the pattern of previous work ^34, 35^, there was a robust effect of the *BDNF* Val^66^Met polymorphism on memory performance. Individuals homozygous for the *BDNF* Met allele demonstrated poorer performance on tests of visual memory relative to those homozygous for the *BDNF* Val allele. Importantly, this was also consistent with the current results of the impact of the *BDNF* polymorphism on visual LTP. LTP magnitude decreased with the increasing number of Met alleles an individual carried. This effect was seen in the *post-hoc* comparisons between all three groups in the late post-tetanus block, and between Val homozygotes and Met carriers in the early post-tetanus block. Spriggs, Sumner et al., ^15^ also found that the Val^66^Met polymorphism impacted the magnitude of visual LTP, with genotype differences in the P2 component of the VEP (greater shift in Met carriers)^i^. It will therefore be important for further studies to characterize the complex relationship between the *BDNF* Val^66^Met polymorphism and visual LTP.

Nevertheless, these data provide compelling support for the hypothesis that differences in memory task performance between BDNF genotypes are, at least to a considerable extent, due to differences in acute or rapid LTP-*like* changes in synaptic transmission in mnemonic networks ^15, 34, 35^. While there may be chronic developmental differences in brain structure resulting from the *BDNF* polymorphism ^33^, here we demonstrate a genetic difference in experience-dependent plasticity that is over and above any developmental differences. It is interesting to note that, in light of relatively high memory scores, the Val/Met group showed little LTP and the Met/Met homozygotes showed on average the inverse of LTP (long-term depression (LTD)). We have previously noted that the baseline stimulation used in the current paradigm can induce LTD in specific groups ^11, 14, 25^, which may be due to modulatory or metaplastic processes (see ^53^ for review). It will therefore be important for future studies to examine the influence of genotype on LTP/LTD thresholds.

It should be noted here that the sample size in this study is small relative to the sample sizes considered ‘adequate’ to yield sufficient power in most behavioural genetics studies. However, as highlighted by Rasch et al., ^54^, sample sizes as low as 20 participants have previously been sufficient to identify genetic effects in imaging studies due to the increased proximity, and thus sensitivity, of neuronal phenotypes to the effect of the polymorphism ^35, 54–56^. The current results support this hypothesis. Specifically, while there was moderate evidence for an effect of the *BDNF* Val^66^Met polymorphism on memory performance, the evidence was overwhelming for the effect of the polymorphism on LTP magnitude. Additionally, there was a dosage effect of genotype on LTP that was only trending for memory scores. While it is important to acknowledge that replication will be critical, the current results do indicate that the effect size of genetic variations on brain activity is much larger than on behavioural measures, rendering a small sample size less pertinent.

Aberrant plasticity is implicated in a number of psychological and neurological conditions, including schizophrenia ^57^ and Alzheimer’s disease ^58^. Previous studies using the sensory LTP paradigm have demonstrated reduced LTP magnitude in both major depression ^26^ and schizophrenia ^27^. As previously mentioned, the memory tasks employed in the current study have also been used widely in clinical assessments of memory function ^19–22^. We have demonstrated for the first time that the combination of these neurophysiological and behavioural measures provides a level of insight beyond the use of these measures in isolation. With a focus on the *BDNF* Val^66^Met polymorphism, the current study demonstrates how this can be used to explore the neural-basis for group differences in memory performance. It is hoped that this will be of utility to future studies assessing memory decline in neuropsychological and neurodegenerative disorders.

Here we provide the first evidence for the relationship between visually-induced LTP and visual memory performance. This not only bridges the gap between LTP and memory performance, but provides further evidence for the visual-LTP paradigm as an effective index of memory related neuroplasticity in the human neocortex. Additionally, the current study demonstrates that both memory performance and LTP magnitude are influenced by the *BDNF* Val^66^Met polymorphism, thus supporting the role of the polymorphism exerting its influence over memory performance through the modulation of experience-dependent plasticity. Finally, the current study demonstrates the unique insight offered through the combination of these neurophysiological and behavioural measures of memory function. It is anticipated that this will have clinical applications in studying the variety of cognitive and affective disorders.

## Acknowledgements

This work was supported by a grant from the NZ Royal Society (Marsden Grant #06-UoA-077; IJK, KEW). MJS is supported by a Brain Research New Zealand Doctoral Scholarship. The authors would like to thank the Auckland Sequenom Facility for genetic analysis. Subsets of these data have been presented at conferences, and have thus been described previously in published abstracts (e.g. Thompson et al., 2010; 2011; 2013).

## Financial Disclosures

There are no conflicts of interest to report

## Footnotes

i While the results of these two studies may seem contradictory, there are significant differences in the approaches taken between them. Firstly, Spriggs, Sumner et al., (2017) grouped Met homozygotes and heterozygotes together into a ‘Met Carrier’ group, thus overshadowing the dosage effect seen in the late post-tetanus block of the current study. Additionally, the current study employed previously established analysis methods and specifically focused on the N1b. Spriggs, Sumner et al., (2017) analyzed the entire time window from 0-250ms as this was better suited for the use of Dynamic Causal Modelling in this study, but also may have washed out the genotype effect on the N1b.

